# Preprint: A robust peptidomics mass spectrometry platform for measuring oxytocin in plasma and serum

**DOI:** 10.1101/042416

**Authors:** Ole Kristian Brandtzaeg, Elin Johnsen, Hanne Roberg-Larsen, Knut Fredrik Seip, Siri Leknes, Elsa Lundanes, Steven Ray Wilson

## Abstract

Current approaches to measuring the cyclic peptide oxytocin in plasma/serum are associated with poor selectivity and/or inadequate sensitivity. We here describe a high performance nano liquid chromatography-mass spectrometry platform for measuring OT in human plasma/serum. The platform is extremely robust, allowing laborious sample clean-up steps to be omitted. OT binds strongly to plasma proteins, but a reduction/alkylation procedure breaks this bond, allowing ample detection of total OT. The method showed excellent quantitation properties, and was used to determine total OT levels to 0.5-1.2 ng/mL (evaluated with human plasma and cord serum). The method is compatible with accessible mass spectrometry instrumentation, finally allowing selective and easily comparable oxytocin measurements.

## Introduction

The neuropeptide oxytocin (OT) is a facilitator of childbirth and breastfeeding, and can activate maternal behavior ^1^ and partner preference ^2^ in animal models. In humans, OT levels have been related to e.g. autism ^3^, and schizophrenia ^4^. Several studies have reported a coordinated release of central and peripheral OT ^5^,^6^ and that peripheral levels can be a low-invasive indicator of central state ^7^. However, the brain/blood OT relation is a disputed topic ^8^ casting doubt on the biomarker-ness of peripheral OT. A key source of skepticism is the absence of satisfactory analytical methodology of OT measurements ^9^. Nearly without exception, enzyme-linked immunosorbent assays (ELISA) and radioimmunoassays (RIA) are used to monitor OT in blood and other biofluids. These methods have in recent years been severely criticized due to poor selectivity ^8^,^9^. An alternative to ELISA/RIA is mass spectrometry (MS). The MS instrument allows unambiguous identification/quantification of e.g. peptides, by first recording the molecular mass of a compound (single MS), and then creating a molecular “fingerprint” by fragmenting the compound to smaller parts (MS/MS). Separating compounds in a mixture (e.g. plasma) prior to MS detection further strengthens identification and sensitivity. Peptides are typically separated using liquid chromatography (LC). LC-MS is an invaluable tool in virtually all areas of biomedical analysis. A notable exception is however OT measurement; the few published methods for LC-MS measurements of plasma OT ^10,11^ provide unsatisfactory sensitivity and varying results, and are therefore difficult to put to practical use. We here set out to develop a robust and sensitive method for quantification of OT in blood, as a remedy for the under-par LC-MS and ELISA/RIA performance regarding OT analysis. We here “borrow” tools from mass spectrometry based proteomics, namely i) nanoLC-MS (a particularly sensitive variant of LC-MS ^12^) featuring on-line sample extraction ^13^, and ii) a reduction/alkylation step ^14^, allowing vastly increased OT extraction.

## Results

### Enabling nanoLC-MSfor robust and simple plasma analysis

NanoLC-MS is exceptionally sensitive and selective instrumentation for identifying and measuring e.g. peptides ^12^, and is commonplace in proteomics facilities. However, nanoLC-MS is rarely used for large-scale blood/serum/plasma sample analysis, in part due to its limited robustness (i.e. clogs easily if extensive sample preparation is not undertaken). This weakness was overcome by implementing an automated filtration/filter back-flush (AFFL) unit ^15^ to the nanoLC-MS system, allowing robust plasma analysis. Details are described below.

In preliminary experiments with a standard nanoLC-MS set-up (i.e. trap column for extraction + separation column), injecting protein precipitated pooled human plasma clogged the column(s) (see **Figure 1 A**). After just one plasma injection, it was not possible to reuse the columns, even after extensive washing attempts. Therefore, we incorporated an AFFL system upstream to the nanoLC-MS platform (see **Figure 1 B-C**). AFFL allows samples to pass through a stainless steel filter that captures particulate matter; this matter is flushed backwards off the filter after each injection, allowing filter intactness (and hence system robustness) for very large numbers of injections ^15^. To illustrate, only a minimal increase in back pressure between the first to the hundredth plasma injection was observed (**Figure 1 A**). OT spiked to plasma could be chromatographed with excellent retention time repeatability (0.1 % RSD; see **Figure 1 D**). During this study, over 300 samples were injected without need for part/column replacement. Taken together, AFFL-SPE-nanoLC-MS is a highly suited platform for blood peptidomics, e.g. targeted determination of oxytocin.

**Figure 1:**
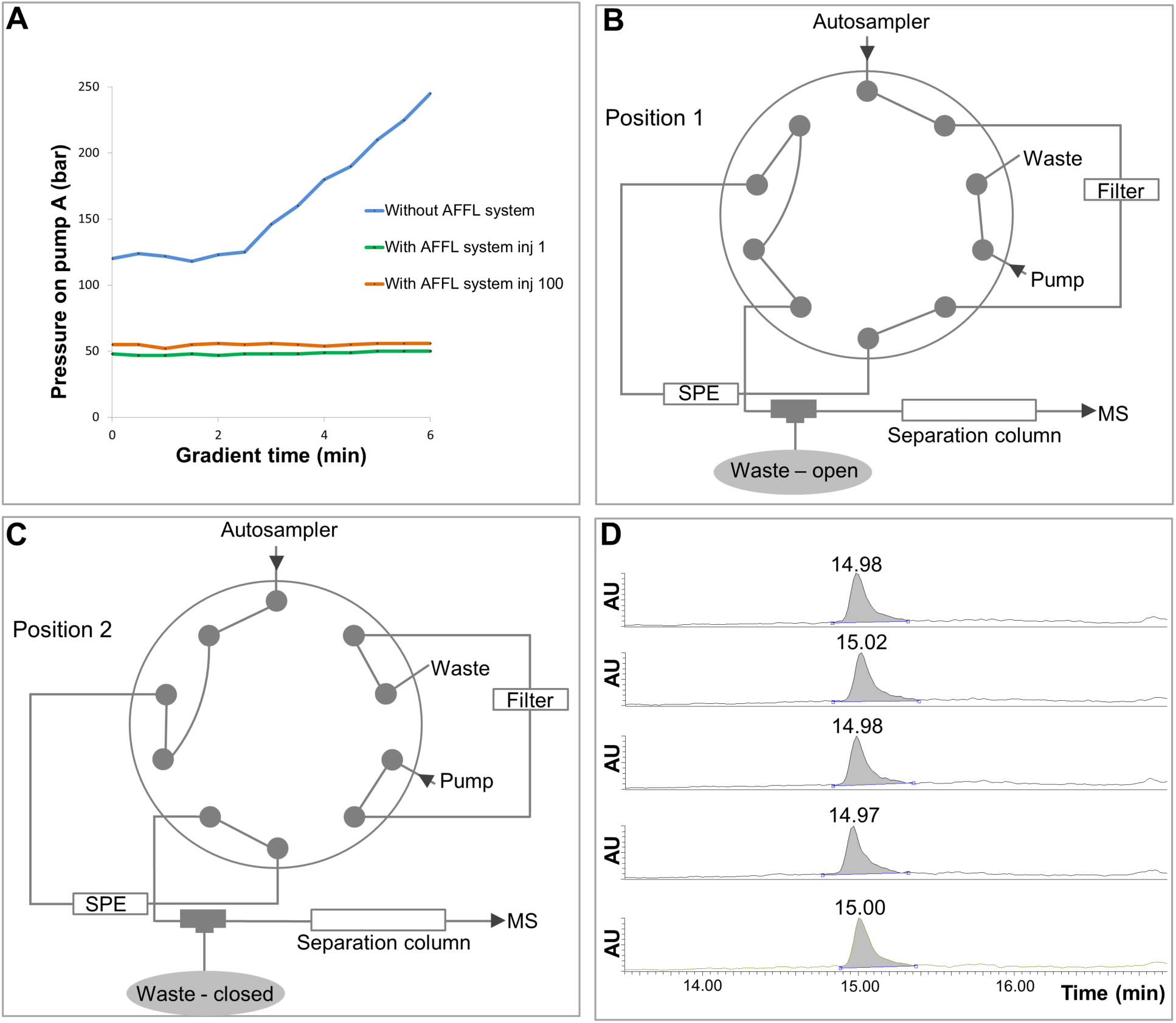
AFFL-SPE-nanoLC-MS for plasma analysis. **A.** Pressure profiles on pump A of the Easy nl_C pump, during a 6 minutes gradient (0-90% B) at a flow rate 800 nL/min, when injecting a plasma sample without the AFFL system (blue line). The green line illustrates the pressure profile during the gradient for the first plasma sample injected when the AFFL system was incorporated, while the orange line illustrates the pressure profile during the gradient for the hundredth plasma injection onto the AFFL system. **B.** Position 1 of the external 10-port valve. In this position the particles are retained on the filter, while hydrophobic compounds (including OT) is retained on the SPE and salts and hydrophilic compounds are eluted to waste. **C.** Position 2 of the external 10-port valve. The filter is being back-flushed, and hydrophobic compounds are eluted off the SPE and separated on the separation column before detection by MS. **D.** Five injections of plasma sample spiked with OT to a final concentration of 500 pg/mL.

### Sensitive and stable detection of plasma OT following a reduction/alkylation step

We find that OT strongly binds to plasma proteins, which can seriously affect the sensitivity/measuring accuracy in biomarker studies. However, performing a reduction and alkylation step liberates OT from plasma proteins, allowing ample sensitivity and precise quantification of endogenous (total) OT. Details are described below.

Initially, samples contained 50 mM ZnCl_2_ (10 mM aspartate buffer, pH 4.5) to stabilize OT via chelation ^16^ prior to subsequent sample preparation (e.g. removing proteins via protein precipitation (PPT)). However, adding ZnCl_2_ to plasma samples resulted in noisy signals and pressure build-up, likely due to on-column precipitation of salts and/or proteins. Acetonitrile based PPT (without the presence of chelating agents) was associated with an unassuring recovery profile (OT recovery dropped and leveled off after 40 minutes (**Figure SM 1**)). OT was stable in the solvents used during and after PPT (**Figure SM 2**), and did not absorb to tubes and vials. It was considered unlikely that the main metabolizing enzyme for OT in plasma, cystinyl aminopeptidase/oxytocinase ^17^ was degrading OT in these conditions, as this enzyme is rather large (subject to PPT), and blood from non-pregnant individuals was used. Therefore, we speculated that the recovery profile depicted a slow binding to protein remains. To further assess the issue of OT protein binding, pooled human plasma was spiked with oxytocin, and was stored on the laboratory bench up to 8 h before PPT; recovery of the spiked OT linearly deteriorated as function of time before the PPT step (**Figure SM 3**), once again suggesting a slow and strong protein binding after spiking. Furthermore, OT spiked to plasma had very poor filtrate recovery using size separation with centrifugal filters, again implying strong protein binding.

We hypothesized that strong protein binding was preventing detection of endogenous OT ((<pg/mL levels, **Figure SM 4**) due to co-precipitation during PPT. The disulfide bridge (DSB) of OT (**Figure 2a**) can engage in complexes ^16^, and likely with serum albumin (buminlbumi, which contains multiple DSBs. To obstruct plasma protein binding, a reduction/alkylation (R/A)^14^ step was performed which irreversibly breaks DSBs (**Figure 2b**). When analyzing unspiked R/A treated plasma samples, endogenous OT was found to be present at strikingly high levels (see LC-MS chromatogram, **Figure 2b** and **Figure SM 5**). OT was determined in pooled plasma and human cord serum, obtained from commercial sources: The concentration of oxytocin in pooled human plasma from Sigma Aldrich and Innovative Research was 0.5 ng/mL and 0.7 ng/mL, respectively. For pooled human cord serum (Innovative Research) the OT concentration was expectedly higher^18^,1.2 ng/mL (**Figure 3a**). Oxytocin plasma levels were, as expected, higher after nasal intake of OT (**Figure 3b**). However, the fold-change was very dependent on the individual. For instance, person 2 (who described him/herself as highly anxious prior to sample collection) had a markedly different OT plasma profile before/after intranasal administration. Our results confirm the common assumption that OT levels can significantly vary between individuals^19^ (Identification/quantification of OTwas based on using external standards, a deuterated internal standard, and characteristic MS/MS transitions for quantification/qualification. The quantitative traits of the assay included excellent linearity (5-2000 pg/mL, r^2^ = 0.999), high recovery (90 %) and good precision/reliability (RSD: 0.4-4.3 %, depending on concentration); see **Figure SM 6**.

**Figure 2:**
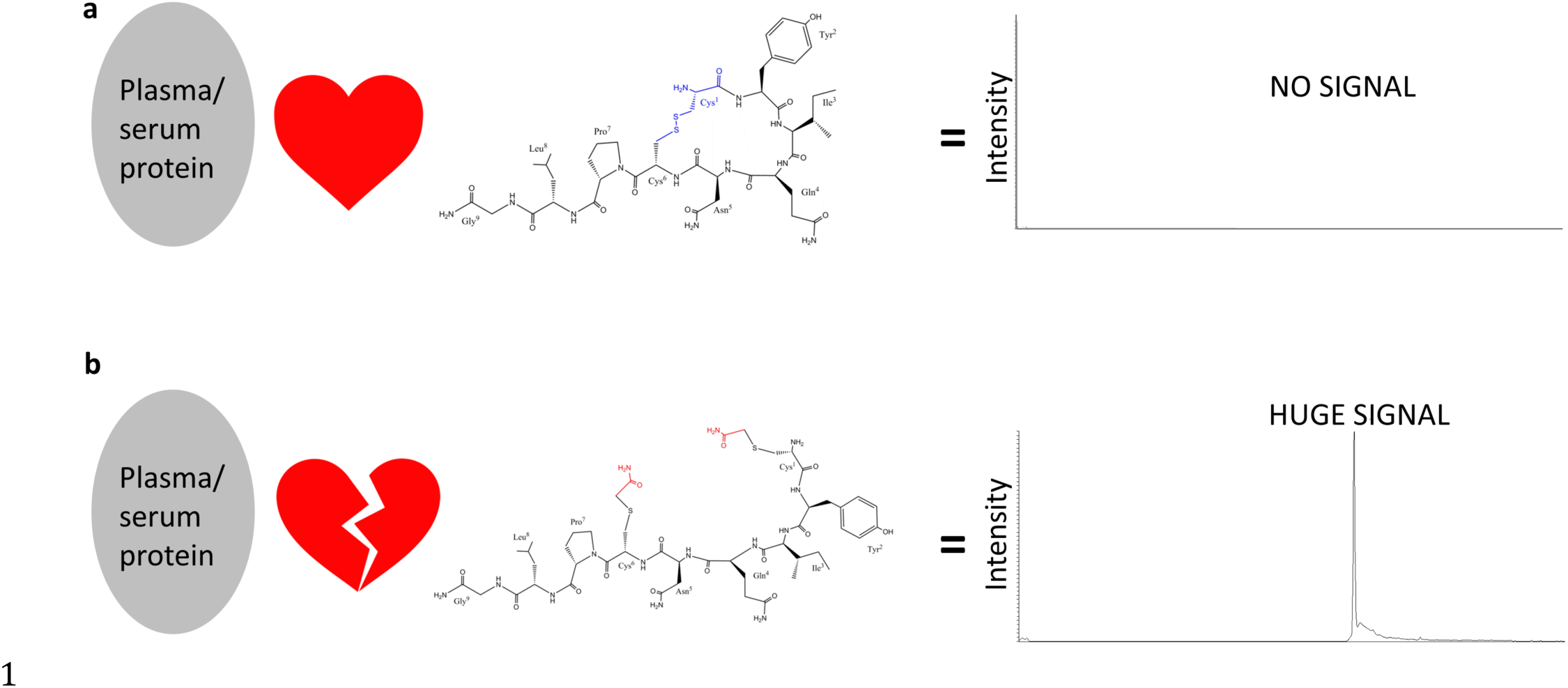
Effect of reducing and alkylation on oxytocin measurement in plasma. **A.** OT binds to plasma/serum proteins, and co-precipitate during sample preparation, resulting in poor detection. **B.** Reduction and alkylation breaks OT binding to plasma/serum proteins, preventing loss of oxytocin during sample preparation, resulting in ample detection.

**Figure 3:**
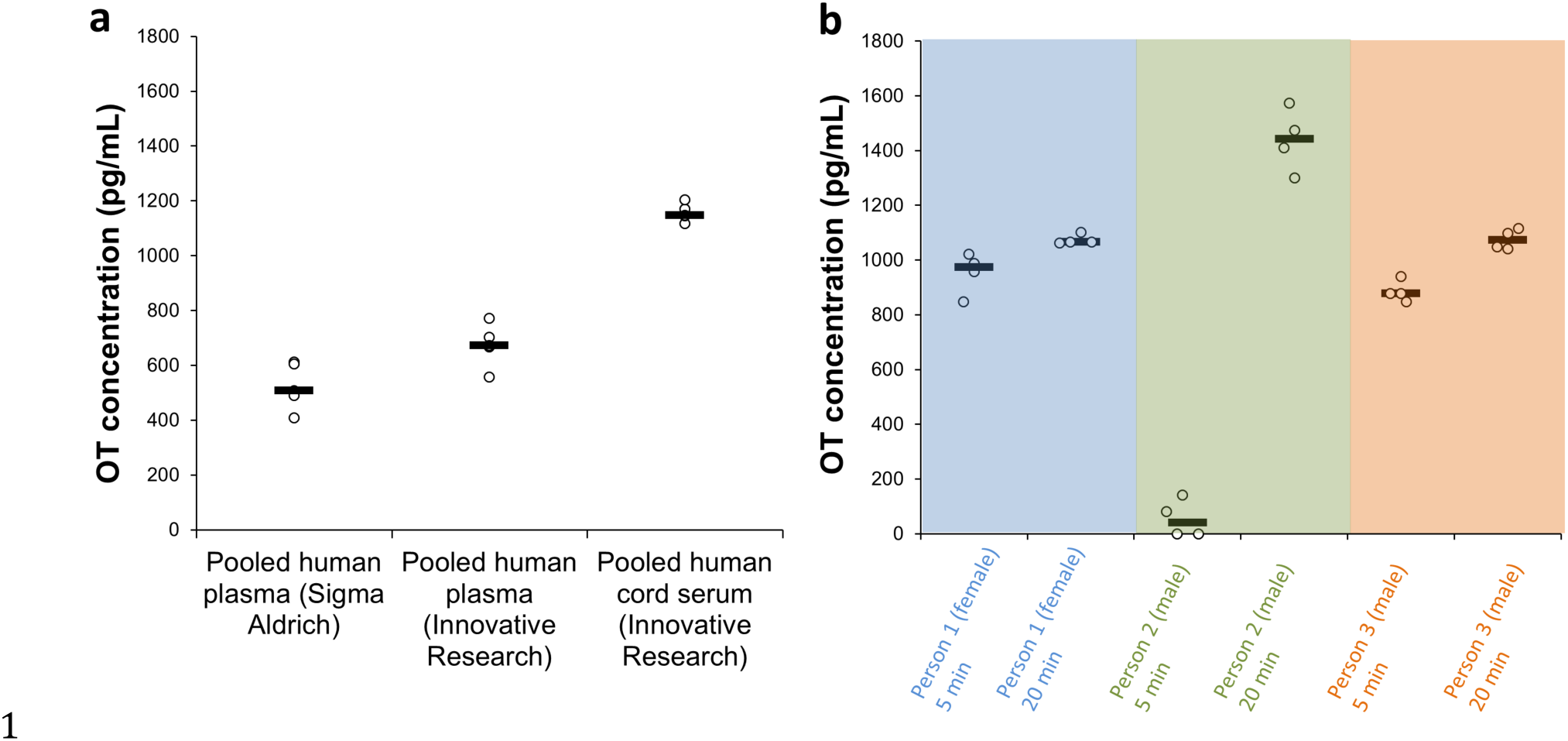
Oxytocin levels in human plasma/serum. **A.** OT basal levels in pooled human plasma from two vendors (Sigma Aldrich and Innovative Research), and OT basal levels in pooled human cord serum from Innovative Research. **B.** OT plasma concentration from three individuals sampled 5 and 20 minutes after applying two puffs of OT nasal spray in each nostril.

## Discussion

A reduction and alkylation step was key in “liberating” oxytocin from plasma proteins, allowing ample detection of endogenous high pg-ng/mL amounts in human plasma. Tight plasma binding is not uncommon with biomarkers ^20^. The OT levels observed in this study are remarkably higher compared to that obtained with an off-line extraction step (low pg/mL levels) ^21^. With extraction, the vast majority of OT is discarded with plasma proteins, leaving only a minute free amount of OT left to be measured. Measuring only the free fraction, as currently recommended (ref leng, mccullogh) can be a confounding factor, since the free OT concentration can be drastically changed by factors such as age, morbidity, or by compounds that displace OT from proteins ^22^. This is especially the case if the marker is heavily bound ^22^, as we find with OT. Indeed, even using MS large variations are observed when measuring the free fraction of OT; a third of the human samples analyzed by Zhang et al did not contain detectable levels of OT ^10^. We have also registered such inconsistencies with our own “neurotransmitter-omics” MS platform ^11^. In addition, free OT levels varied 6-fold within a homogenous group of rats ^10^. As shown in Figure 3, when all circulating OT is measured using our method there differences between individuals are already pronounced individuals (but not unusually large compared to much of the metabolome). Such individual differences are thought to be highly informative ^19,23^; additional confounding factors will undoubtedly make correlations less clear. Based on this reasoning, total OT is better suited as a biomarker than only the free fraction of OT.

Considering the growing concerns of antibody-based assays (both RIA and ELISA fall in to this category) regarding selectivity and antibody kit reproducibility ^24^, LC-MS is a natural choice for OT measurements due to its excellent selectivity. The robust and highly automated AFFL-nanoLC-MS approach has attractive quantification traits, and can be simply implemented in e.g. proteomics facilities (common in e.g. many larger universities/hospitals). As the instrumentation is compatible with salty solutions, AFFL-nanoLC-MS can be used for urine and cerebral spinal fluid measurements as well (protein binding can also occur in these matrices). Other LC-MS systems can be employed, e.g. UPLC-MS systems used for drug measurements or metabolomics, but these may require off-line filtration/extraction steps.

## Methods

### Chemicals and reagents

Oxytocin (OT) acetate salt hydrate (?97 %), oxytocin-d_5_ (9 8%, internal standard (IS)), dithiothreitol (DTT), iodoacetamide (IAM), acetonitrile (LC-MS grade), formic acid (FA, LC-MS grade) and pooled human plasma with 4 % trisodium citrate as anticoagulant (P9523-5mL, lot#: SLBK0464V) were purchased from Sigma Aldrich (St. Louis, MO, USA). Pooled human plasma with EDTA as anticoagulant (lot#: 17964) and pooled human cord serum (lot#: 18241) were obtained from Innovated Research (Huntsville, AL, USA). 1 M Tris-HCl pH 8.0 was made by Oslo University Hospital (Oslo, Norway). LC-MS grade water was bought from Fischer Scientific (Hampton, NH, USA), while type 1 water was acquired from a Milli-Q^®^1 Integral 5 water purification system (Merck Millipore, Billerica, MA, USA).

### Storage of stock solutions, plasma and serum

Stock solutions of OT (5 μg/mL) and IS (10 μg/mL) dissolved in LC-MS grade water, pooled human plasma and pooled human cord serum were stored in freezer at −20o C.

### Preparation of calibration standards and samples

For all standard solutions and plasma/serum samples, 10 μL of a 10 ng/mL working solution of IS were added so that the concentration in the final reconstitution volume (100 μL) was 1 ng/mL IS. All solutions were made in 1.5 mL Eppendorf LoBind tubes (Hamburg, Germany). Standard solutions used for establishing the calibration curve were made by appropriate diluting a working solution of 10 ng/mL OT in 0.1 % FA with 0.1 % FA to a final concentration in the reconstituted solutions of 5, 500, 1000 and 2000 pg/mL. Dilution of the plasma/serum samples and standard solutions was performed by pipetting (with newly calibrated pipettes) 100 μL of plasma/serum samples and standard solutions into 200 μL 50 mM tris-HCl (pH 8.0). For reduction of disulfide bonds, 5 μL of 0.5 M DTT were added to all solutions followed by whirl mixing for 30 sec, incubation at 37°C for 45 min, and finally cooling to room temperature (22°C). Alkylation was done by adding 15 μL of 0.5 M IAM into each solution followed by whirl mixing for 30 sec before incubation at 22°C in the dark for 20 min. Protein precipitation was performed by adding ice-cold 80 % ACN in LC-MS grade water (v/v), and whirl mixing for 30 sec before centrifugation for 15 min at 14,000 relative centrifugal force (rcf) in an Eppendorf 5415 R-model centrifuge (20oC) (Hamburg, Germany). The supernatant was pipetted into a new tube and evaporated to dryness in a Speed Vac^®^1 SC110-model from Savant, Thermo Fisher Scientific (Waltham, MA, USA), followed by reconstitution in 100 μL 0.1% FA in LC-MS grade water (v/v). Aliquots of 10 μL of this solution were analyzed by the nanoLC-MS/MS platform.

For investigating protein binding, OT was spiked into human plasma and 500 μL was applied to 10K Amicon^®^1 ultra centrifugal filters from Merck Millipore (Billerica, MA, USA). An aliquot of 20 μL of the filtrate was analyzed by the Bruker Easy nLC system (without AFFL) connected to a TSQ QuantivaTM triple quadrupole mass spectrometer from Thermo Scientific.

### Nasal spray experiment

Three healthy volunteers, one female and two males were asked to apply two puffs of OT nasal spray (6.7 μg OT/puff, Syntocinon^®^1 from Sigma-Tau Pharmaceuticals, inc., Gaithersburg, MD, USA) in each nostril. The subjects were asked to spray close to the respiratory region, where there has previously been shown best absorption ^25^. Two blood samples were drawn from each participant; one 5 min and another 20 min after the puffs of OT nasal spray were applied. Plasma was made and the samples were analyzed the same day (n=4).

### Automatic filtration and filter back-flush (AFFL) solid phase extraction nanoLC tandem MS peptidomics platform

An EASY-nLC liquid chromatograph with an integrated 6x4 autosampler from Bruker (Billerica, MA, USA) was used as pump. Mobile phase A was 0.1% FA in LC-MS grade water (v/v), while Mobile phase B was 0.1% FA in LC-MS grade acetonitrile (ACN). The loading mobile phase composition was 0.1% FA in LC-MS grade water. The external 10-port valve from VICI (Schenkon, Switzerland) controlled by the MS-software was used in the AFFL system. See Figure 1 for plumbing of the AFFL system. A Hitachi L-7100 HPLC pump (Chiyoda, Tokyo, Japan) in isocratic mode was used to back-flush the filter in the AFFL system with type 1 water. In position 1 (Figure 1 A), the sample passed through a stainless steel filter (1 μm porosity, 1/16” screen, VICI) onto a 100 μm ID x 50 mm silica monolithic C18 SPE manufactured as described in ^26^ (similar to Chromolith^®^1 CapRod C18 capillary columns from Merck Millipore). In position 2 (Figure 1 B), two processes happened simultaneously; the filter is back-flushed, while oxytocin is back-flushed from the SPE column onto a 100 μm ID x 150 mm silica monolithic C18 separation column manufactured as described in ^26^ (similar to Chromolith^®^1 CapRod C18 capillary columns from Merck Millipore). A steel emitter, 30 μm ID x 40 mm, from Thermo Scientific, was connected to the end of the separation column by a 1/16” standard steel internal union from VICI. A nanospray Flex^TM^ ion source (nanoESI) coupled to a Quantiva^TM^ triple quadrupole mass spectrometer from Thermo Scientific was used for detection of oxytocin in full MS-and tandem MS-mode (MS/MS).

### Liquid chromatography and mass spectrometry parameters

The 20 min gradient program was composed as follows: 20 %B isocratic elution for 14 min, followed by an increase from 20 to 90 % B in 2 min before isocratic elution at 90 % B for 4 min. The injection volume was 10 μL. The SPE was equilibrated with 4 μL 0.1 % FA in LC-MS grade water at a constant flow rate of 3 μL/min, while the separation column was equilibrated with 5 μL 0.1 % FA in LC-MS grade water at a flow rate of 3 μL/min before each injection. The MS was operated in positive MS-mode and selected reaction monitoring (SRM) mode was used. The spray voltage was set to 1.6 kV. The precursor ions for native oxytocin and IS were *m/z* 1007.475 and *m/z* 1012.475, respectively. For oxytocin the product ions were *m/z* 285.125 with 38 V collision energy (CE), and *m/z* 723.225 with 30 V CE. For IS the product ions were *m/z* 290.125 with 38 V CE, and *m/z* 723.225 with 30 V CE. The precursor ions for reduced and alkylated (R/A) oxytocin and IS were *m/z* 1123.547 and *m/z* 1128.547, respectively. For R/A oxytocin the product ions were *m/z* 285.125 with 38 V CE, and *m/z* 839.302 with 30 V CE. For R/A IS the product ions were *m/z* 290.125 with 38 V CE and *m/z* 839.302 with 30 V CE. The Q1 and Q3 resolutions were both set to 1.2 FWHM, and the RF lens had a voltage of 185. A cycle time of 1 sec was used with 3 mTorr collision-induced dissociation (CID) gas. Argon was used as collision gas. In addition, 25 V source fragmentation energy was used together with 3 secs chrom filter.

### Data analysis and interpretation

Data analysis and interpretation were done using Xcalibur^TM^ software version 3.0 from Thermo Scientific.

### Ethical statement

All subjects gave written informed consent, and the blood collection was approved by the Regional Ethics Committee (2011/1337/REK S-OE D). All methods were carried out in accordance with the approved guidelines and regulations.

